# Affective arousal explains infant gaze following under various social context

**DOI:** 10.1101/2022.02.04.479127

**Authors:** Mitsuhiko Ishikawa, Atsushi Senju, Masaharu Kato, Shoji Itakura

**Affiliations:** Centre for Baby Science, Doshisha University, 4-1-1 Kizugawadai, Kizugawa, Kyoto 619-0295 Japan; Centre for Brain and Cognitive Development, Birkbeck, University of London, Malet Street, London WC1E 7HX, UK; Research Centre for Child Mental Development, Hamamatsu University School of Medicine, 1-20-1 Handayama, Higashi-ku, Hamamatsu, Sizuoka 431-3192 Japan

**Keywords:** Eye Contact, Reliability, Gaze Following, Heart Rate, Social Decision Making

## Abstract

Gaze following is fundamental to human sociocognitive development, such as language and cultural learning. Previous studies have revealed that infant gaze following is not a reflexive orienting to adult’s eye movement. Instead, infants adaptively modulate gaze following behaviour depending on social contexts. However, the neurophysiological mechanisms underlying contextual modulation of gaze following remain unclear. We tested whether contextual modulation of infant gaze following is mediated by the infants’ heart rate, which is hypothesised to indicate the calculation of action value. Forty-one 6- to 9-month-old infants participated in this study. Infants observed either a reliable face, which gazed toward the location of an object, or an unreliable face, which gazed away from the location of an object. Then, the infants watched a video of the same model making eye contact or not showing any ostensive signals, before shifting her gaze toward one of two objects. We revealed that reliability and eye contact independently increased heart rates, which then fully mediate the effect of these social cues on the frequency of infant gaze following. Results suggest that each social cue independently enhances physiological arousal which then accumulatively predicts the likelihood of infant gaze following behaviour.

## Introduction

A hallmark of human intelligence is an exceptional capacity to learn from other humans [1, 2]. From the earliest stage of life, learning from other conspecifics is essential for human infants to obtain information relevant to their survival and acculturalisation, including the acquisition of generalisable knowledge or skills from another individual [3]. Especially in preverbal infants, adult’s gaze direction plays a critical role in social learning [4]. Thus, it is not surprising that infant gaze following (GF) behaviour has become the core topic in cognitive, developmental, social, and comparative psychologies and neurosciences [5].

Early studies saw infant GF as an automatic or reflexive shift of infant visual attention, induced by adult’s gaze direction [6]. However, studies in the past decade started to reveal that infant GF is context-dependent, in which various social contexts modulate the probability of infant GF. For example, infants follow others’ gaze when accompanied by ostensive signals, the signal which is hypothesised to convey an adult’s communicative intent [7, 8]. Senju and Csibra [7] showed that 6.5-month-old infants follow others’ gaze when it followed eye contact or infant-directed speech (ostensive signals), but not when it followed non-ostensive but attention-grabbing stimuli (e.g., non-social animation overlaid on top of the actor’s face). Csibra & Gergely [3] interpreted this (and convergent) evidence that infants follow others’ gaze when they refer to the topic of communication within the framework of ostensive-referential communication, and suggested the theory of natural pedagogy.

The trait of the communicative partner, such as reliability as an informant, also modulates infant GF and subsequent selective learning [9, 10]. It is reported that 8-month-old infants can track the reliability of potential informants and use this information judiciously to modify their future behaviour [11]. It has been shown that infants modulate their social behaviour such as imitation [12, 13] and GF [14] depending on others’ reliability. In two of these studies [12, 14], infants first observed an experimenter looking inside a container that either contained a toy (reliable looker condition) or was empty (unreliable looker condition). In this task, infants could expect to find a toy when the experimenter gazes inside a container or to find the container empty. After the observation, infants engaged in the GF task [14], thus it was possible to test whether infants’ GF would be modulated by their knowledge of the experimenter’s gaze reliability. This kind of observing experience of other’s gaze behaviour made it possible for the infants to evaluate other’s reliability as an informant, and the infants thereby modulated their social engagement. Other factors such as emotional states of the model signalled by her facial expression [15] and perceptual status of the model (e.g. whether she can see the target object or not) [16] are also known to modulate infant GF behaviour.

The studies demonstrated that infants are more likely to follow gaze when they are relevant or informative. It has been suggested that ostensive signals activate the presumption of relevance in infant learners [17, 18]. Also, it has been shown that infants have sensitivity to the reliability of informants, and infants can keep their knowledge to track the reliability of potential informants [19]. These different contextual modulations of GF may have common cognitive substrates before emerging as behaviours. We hypothesise that infants calculate ‘action value’ in social context from various contextual cues, which then increase the probability of infant GF if infants expect reward based on the value calculation. In this framework of value calculation, infants are expected to select the action which has a higher value than alternative actions. The frameworks also assume that infants expect reward following the action execution, which is again based on the value calculations. Reward-driven learning is a fundamental learning form from early infancy [20] and suggested as a principal mechanism of learning [21]. Triesch et al. [22] applied reinforcement learning to the computational modelling of the emergence of GF, and suggested that infants’ GF is reinforced by rewarding experiences in the mother-infant interaction and infants select GF according to high action value. Also, Deák et al. [23] argued that salient and strong reward signals may enhance the frequency of GF behaviour. We hypothesise that infants calculate action values in social contexts, and the calculated value for each action (e.g. to follow an adult’s gaze), or expected reward following the execution of such action, influence infant’s decision to follow (or not to follow) gaze. From this perspective of action value calculation, different contextual cues are processed and integrated as an outcome of value calculations, then the value of which will determine the modulation of GF behaviour.

Because the reward system is deep inside of human brains, it is technically difficult to measure brain activations directly in infants, particularly for GF behaviour which requires visual processing of fully awake infants during imaging [24]. To overcome this challenge, recent studies from our lab monitored infant physiological arousal, as indicated in the change in heart rate (HR), during the GF situation. The assumption that physiological arousal is indicative of the processing of the reward system is derived from previous studies, which suggest that physiological arousal is indicative of reward expectation [25, 26, 27]. For example, a recent study showed that reward predictive cues can enhance physiological arousal in 7-month-old infants [25]. Furthermore, infants in this study chose to predictively look at the place where a rewarding animation will be presented, suggesting that reward expectation can modulate infant gaze behaviour. Thus, although indirect and somewhat speculative, we propose that HR measurement is one of the most promising ways to explore infant action value calculation in social contexts. Supporting this position, our recent study examined infant physiological arousal with HR measurement, the result of which suggested that physiological arousal, which we hypothesise to indicate reward expectation, mediate contextual modulation of infant GF. The study showed that eye contact elevated heart rate in infants, and that heart rate increase predicted later GF [28]. In other words, it was suggested that an increase of physiological arousal mediate the subsequent GF behaviour in such an experimental context. This is particularly relevant and necessary to better understand the contextual modulation of infant GF, because some behavioural studies have reported that infants show GF without ostensive signals [29, 30], suggesting that eye contact (or other ostensive signals) does not automatically determine the subsequent GF behaviour and possible context-specific intermediatory processes are in place. Measurements of physiological arousal in the GF situation could provide an index of possible modulatory processes, which we hypothesise as action value calculation, and help us better understand the mechanisms underlying the execution of infant GF.

However, the results we introduced above does not fully support our hypothesis that physiological arousal is indicative of calculated action value, which integrates multiple socially relevant (or reward predictive) cues. It could still mean that physiological arousal mediates the influence of a specific type of social context, the presence of an ostensive signal, on infant gaze following. To test this hypothesis further, it is essential to show that the increase of physiological arousal mediates the heightened probability of infant GF in other social contexts, beyond the presence or absence of ostensive signals. In other words, it is necessary to measure physiological states in GF situations with other contextual cues which would also be hypothesised to induce reward expectations, and demonstrate that such other contextual cues would increase infant HR, which then mediate the increase in infant GF behaviour.

As we introduced earlier, it has been shown that the reliability, just like the presence of eye contact, facilitates infants’ GF [7, 14, 28]. However, these two contextual factors are qualitatively different: The presence of eye contact is an immediate visual context, while the factor of reliability is knowledge learned from prior experiences requiring memory modulation. Most critically, it has not been tested whether infants’ knowledge of the reliability of social partners affects physiological arousal, which then mediates infant GF behaviour. By empirically examining how physiological arousal and GF behaviour are modulated by the knowledge of the reliability of the social partner and contrasting them against the modulation by perceived eye contact, it is possible to discuss whether these different contextual factors are integrated into physiological arousal and affect infants’ social behaviour, as we hypothesised based on action value calculation.

To test the prediction based on the hypothesis that infants integrate multiple social cues based on action value calculation, which is indicated by infant physiological arousal and explain the subsequent infant GF behaviour, the current study examined how the infant’s heart rate changes in two separate social contexts which have been reported to enhance infant GF, the presence of ostensive signals (communication predictive cues) and looker’s reliability (a learned predictiveness or learned value of others, Heyes [10]). To manipulate a looker’s reliability as an informant, we first showed gaze-cueing situations, in which a female directly gazed toward an object or gazed away from it. Infants could learn that they can acquire rewards by following the female gaze. Then, the same female appeared in a GF task with or without eye contact. Replicating the previous study [28], we predicted that infants would show facilitated GF after eye contact, which would be mediated by increased heart rates. Also, we predicted that if the looker gazed toward an object in the gaze-cueing situation (reliable informant), infants would show more frequent GF toward the same looker, which is also mediated by increased heart rates. If infants’ GF behaviour is determined by the reward expectation indexed by heart rates, which we hypothesise as an index of action value calculations under a certain social context, these two signals independently modulate infants’ heart rate, which would then fully mediate the effects of both social cues (reliability and eye contact) on GF. We also explored whether the effect of reliability and eye contact on GF were additive (i.e. two factors summate to predict the probability of GF) or disjunctive (i.e. presence of either signal is sufficient to maximise GF, with no additive effect of the other signal).

## Methods

### Participants

The final sample for analysis consisted of forty-one 6- to 9-month-old infants completed the study (mean age, 237 days old; range, 180-295 days old), with 21 infants in the reliable looker condition (11 female, 10 male) and 20 in the unreliable looker condition (8 female, 12 male). Previous studies have reported that infants from six months old showed GF in screen-based experimental settings [7, 31], and that there was no difference in GF frequency between six- and nine-month-old infants [32]. The sample size was determined based on a study examining the effects of a looker’s reliability on infant GF [14]. Using the effect size from the previous study (f = .71), we conducted a priori power analysis with G*Power [33]. The result indicated that with 20 participants per group we would have achieved above 95% power with alpha at .05 to find the effects of reliability on infants’ GF. The estimated sample size was also sufficient to find the effects of eye contact on infants’ heart rates.

### Apparatus

We used a Tobii Spectrum Eye Tracker (Tobii pro Lab 1.118, Tobii Technology, Stockholm, Sweden) to record eye movements during the presentation of the stimuli. Sampling rate was 120 Hz. Participants were seated approximately 60 centimetres from the monitor in the caregiver’s lap. Before the recording of infants’ eye movements, a nine-point calibration was conducted.

For the HR recording, we used a BIOPAC MP160 (Biopac System, CA, USA) and a BioNomadix (BIOPAC Systems, CA, USA) with a 3-lead ECG to measure the ECG data at a 1000 Hz sampling rate. Before attaching the electrodes, we used rubbing alcohol to reduce impedance.

### Stimuli and Procedure

Infants participated in two blocks of tasks. Each block consisted of observation of 4 trials of gaze-cueing situations (i.e. induction of one of the reliability conditions), followed by 6 trials of the GF task. This sequence of a block was repeated twice for each participant, which resulted in each infant observing eight trials of the gaze-cueing situations (4 trials x 2 blocks) and performing 12 trials in the GF task (6 trials x 2 blocks). Both blocks were identical per participant. Completing all the trials takes about seven minutes.

### Observation of Gaze-cueing Situations

To manipulate the looker’s social characteristics as an informant, we had infants observe gaze-cueing situations before the GF task. We modified procedures and stimuli used by Ishikawa, Yoshimura, Sato, and Itakura [34]. There were three types of faces: (1) a direct gaze face for a pre-cueing stimulus, (2) a right-gazing face, and (3) a left-gazing face. A target object was placed approximately 15° to the left or right of the centre of the screen per trial. We used four female faces and six colourful illustrations for stimuli. Infants viewed four trials for each block. Each trial began with a picture of a female directly gazing toward the infant (2s), and after that, the female shifted her gaze direction either right or left (1s), with no objects in either direction. Finally, in the third presentation, one of the objects appeared in a place either congruent (reliable looker) or incongruent (unreliable looker) with the female gaze direction (3s). Two reliability conditions were allocated as a between-subject factor, with half the infants only observed a reliable model and the other half only observed an unreliable model. Target objects were randomly chosen from six colourful pictures, such as an apple and a ball. The direction of the model’s gaze was counterbalanced in ABBA order. Half of the infants saw a leftward gaze in the first trial, and the other half saw a rightward gaze first. In a block, the GF task was continuously started after four trial observations of gaze-cueing situations.

### Gaze following task

For the GF task, we applied our stimuli and procedures used by Ishikawa and Itakura [28]. The infants watched a video of the same model who appeared in the gaze cueing situations making eye contact or not showing any ostensive signals, before shifting her gaze toward one of two objects. Each video clip began with a female gazing downward seated at a table. Two toys that were not presented in the gaze cueing situation were placed one at each side of the model. These two toys were alternately assigned as a distractor object and a target object and. The stimuli consisted of three phases. In the Baseline, the model kept still for two seconds and gradually looked up with both eyes closed. After the model facing the front, the Action phase was started, which was different among conditions. In the Eye Contact (EC) condition, the model opened her eyes and kept her direct gaze for three seconds. In the No Cue (NC) condition, the model kept her eyes closed for three seconds. In the Shivering (SV) condition, the model horizontally shook her head a few times for three seconds with keeping her eyes closed. The third phase was the Gazing phase. In the gazing phase, the model directed her head about 45° toward the target object and gazed on it for five seconds. The model opened her eyes just before turning her head in the NC and SV conditions. The model kept her face with neutral expression and make no sounds throughout the entire video clip.

The three communicative cue conditions were designed to run within-subject. Six trials were presented totally to each infant in one block. The order of trials was quasi-randomized across conditions. The assignment of the target object was randomized in a block. The model’s gaze direction was counterbalanced in ABBABA order. Half of the infants watched a leftward gaze in the first trial, and the other half saw a rightward gaze first. The infant’s attention was grabbed to the centre of the screen by a Tobii attention getter before the start of each trial, on which the model’s face appeared in the video clip.

### Data Analysis

The recorded sample’s average percentage of gaze position data through the whole recording was 68.73% (SD = 12.95%, range: 45%-96%). A Clearview fixation filter (Tobii Technology, Sweden) was applied for the eye-tracking data, which is one of the basic algorithms used in the Tobii eye-tracking software and has been used in infant studies [28]. Same as the previous study [28], fixation was defined as gaze recorded within a 50-pixel diameter for a minimum of 200 milliseconds, and this criterion was applied to the raw eye-tracking data to determine the duration of any fixation.

### Eye-tracking data

The measurement of GF was whether the infant’s first fixation toward an object preceded immediately by fixation in the face AOI during the Gazing phase (i.e., after the head turn started) went to the AOI of the object looked at by the model or toward of the object opposite one. If the infant fixated on one of the objects from the start of the Gazing phase, it was not counted as a looking behaviour neither following nor not following. For example, if the infant fixated: (fixation 1) distractor, (fixation 2) distractor, (fixation 3) head, (fixation 4) target), the sequence of the fixation 3 and 4 was counted as gaze following. We followed the same criteria were used in the previous eye-tracking studies of infants’ GF [28, 30], and the likelihood of GF is 50% given this definition of GF. The face AOI includes the top of the head to the chin. The infant’s looking behaviour was coded as GF if the infant fixated at the same object the model gazed at. To be included in the analyses, it was required that infants elicited transition of fixations from the head toward an object during the Gazing phase in at least three trials. Four infants who had fewer than three trials in looking at one of the two objects were excluded from the analysis.

Trials that did not include any fixations from the head toward an object during the Gazing phase were excluded from the analyses. In total, 11 trials were excluded from the analysis of GF (EC: 3 trials, NC: 5 trials, SV: 3 trials).

Also, we analysed the total duration of fixations at the model’s face for the Baseline, the Action, and the Gazing phases separately.

### Heart Rate

We pre-processed the ECG signal using band-pass filtering with 0.5 and 40 Hz cut-off frequencies. Trials with excessive movement of the infant were excluded for the analysis before the calculation of R-wave-to-R-wave (R-R) intervals. Eleven trials were excluded from the analysis of HR. All excluded trials were overlapped with the exclusion of eye-tracking data. In these excluded trials, infants were inattentive, fussy and, moving thus gaze positions and ECG signals could not be measured properly. R peaks were detected by the detection algorithm of AcqKnowledge 3.9.0 software (Biopac Systems Inc., Santa Barbara, CA). Then, R-R intervals were visually checked to find missed beats, which were interpolated with neighbouring R-R intervals. For the interpolation, we added average durations of neighbouring R-R intervals to the missed beats [35]. The ECG data was separated to each of the three phases (Baseline, Action, and Gazing), and the average R-R intervals were calculated in each phase. Beats per minute were calculated for the analysis of HR. The change of average HRs from the Baseline to the Action phase for each trial was calculated to examine whether HR increase predicts GF.

## Results

Data sets used for the analysis are available in the supplement.

### Gaze following

For the analysis of GF, we used the GF rate for each condition, reliability of looker (reliable, unreliable), and communicative cue condition (EC, NC, SV) as independent variables. We conducted a 2 × 3 ANOVA with GF rate, and results showed a significant main effect of communicative cue condition (*F*(2 38) = 6.116, *p* =.003, *η_p_^2^* =. 136). We used Bonferroni correction for all post hoc t-tests. The EC condition showed a higher GF rate relative to the NC and SV conditions (EC vs. NC: *p* =.015; EC vs. SV: *p* =.014).

Also, there was a main effect of reliability (*F*(1, 39) = 5.76, *p* =.021, *η_p_^2^* =.129), and the reliable group showed more frequent GF than the unreliable group. There was no interaction effect between communicative cue and reliability (*F*2, 78) = 1.132, *p* =.328, *η_p_^2^* =.028).

In addition, to examine whether the communicative cue and reliability facilitate GF, we conducted two-tailed t-tests between the GF rates and a chance level of 50%. (see Figure 1) In the reliable group, infants showed significant GF in each communicative cue condition (EC: M = 67.43%, *t*(20) = 3.313, *p* =.004, *d* = 0.74.; NC: M = 59.86 %, *t*(20) = 2.225, *p* =.038, *d* = 0.49; SV: M = 61.1 %, *t*(20) = 2.564, *p* =.019, *d* = 0.57). In contrast, in the unreliable group, infants exhibited significant GF behaviour only in the EC condition (EC: M = 64.55%, *t*(19) = 2.966, *p* =.008, *d* = 0.66.; NC: M = 47.5 %, *t*(19) = −.623, *p* =.541, *d* = −.139; SV: M = 47.05 %, *t*(19) = −.689, *p* =.499, *d* = −.154).

**Figure 1.**
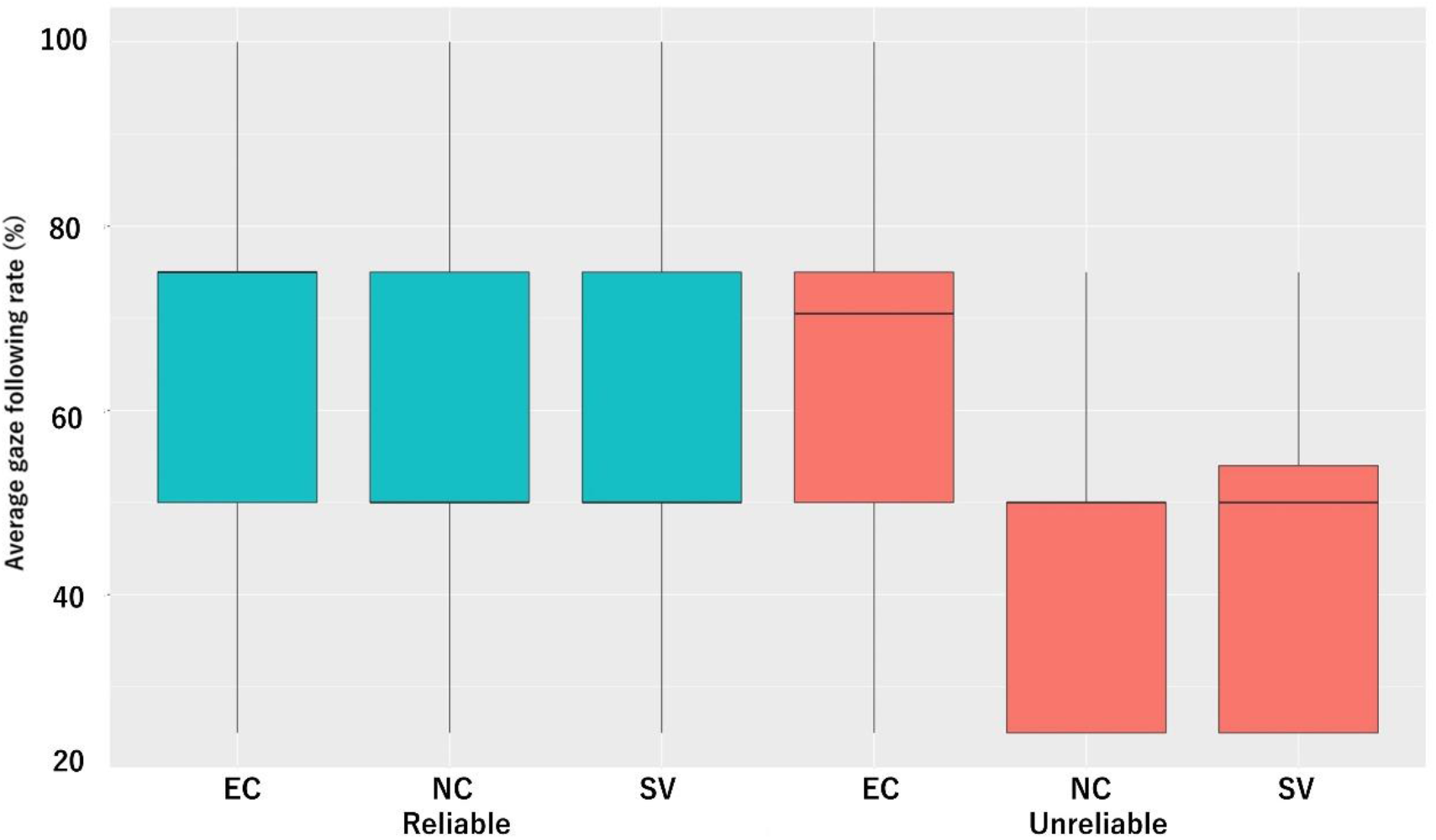
Results of gaze following during the Gazing phase and the proportion of gaze following in each condition. The x-axis depicts condition and the y-axis depicts the percentage of gaze following.

### Heart rate

To examine how infants’ HR changed during the GF situaions, a 2×3×3 ANOVA with two levels of reliability (reliable, unreliable), three levels of condition (EC, NC, SV), and three-phase levels (Baseline, Action, Gazing) was conducted (See Figure S1 in the Supplemental Materials.).

Results showed a significant interaction effect between communicative cue and phase (*F*(4, 156) = 4.535, *p* =.002, *η_p_^2^* =.104). All post hoc t-tests were corrected by Bonferroni correction for multiple comparisons. The post hoc tests showed that HR increase from the Baseline (*M*= 132.11 bpm) to the Action phase (*M* = 132.89 bpm, *p* <.001) and HR decrease from the Action phase to the Gazing phase (*M* = 132.42 bpm, *p* =.001) were found in the EC condition. The SV condition showed a decreased HR from the Action phase (*M*= 132.37 bpm) to the Gazing phase (*M* = 132.09 bpm, *p* =.006). In the Action phase, the average HR was higher for the EC condition than the NC and SV conditions (EC vs. NC: *p* =.032; EC vs. SV: *p* =.039).

In addition, a significant interaction effect between reliability and phase was found (*F*(2, 78) = 4.918, *p* =.01, *η_p_^2^* =.112). The reliable group showed an increased HR from the Baseline (*M*= 132.01 bpm) to the Action phase (*M*= 132.68 bpm, *p* <.001) and a decreased HR from the Action phase to the Gazing phase (*M*= 132.16 bpm, *p* <.001). There was no interaction effect across communicative cue, reliability, and phase (*F*(4, 156) = 0.062, *p* =.993, *η_p_^2^* =.002).

### Predicting gaze following by heart rate increase

The behavioural results found that reliability and communicative cue condition respectively enhanced GF rate. Also, results of HR showed that reliability and communicative cue condition respectively enhanced average HR levels. To examine how contextual factors (reliability and communicative cue condition) modulate HR levels and induce GF, we performed generalized linear model (GLM) logistic regression analyses to predict GF by three factors: reliability, communicative cue condition, and HR increase levels (the degree of he HR increase from the Baseline to the Action phase). The logistic regression of GLM enables us to investigate the influence of factors on the binary response (i.e., following or not) [36], thus it is possible to test whether the HR increase predicts the emergence of GF in each trial. The results of GLM analysis revealed that the HR increase rate predicted later GF (estimate ± SE = 76.45 ± 11.69, *Z*= 6.54, *p* <.001) in all conditions, with higher HR increase pointing to GF behaviour. We also tested the difference between slopes predicting GF by HR increases across the two levels of reliability (reliable, unreliable) and three levels of condition (EC, NC, SV). Results indicated that there were no significant differences between any conditions, in other words, the infants would show GF with the same frequency when they have the same levels of HR increase from the Baseline to the Action phase in any conditions.

### Mediation of contextual modulation on GF by the heart rate increase

The previous GLM analysis showed that the HR increase levels predict the emergence of GF. To examine whether HR increase levels mediate the effects of contextual factors on GF, we compared two models with and without HR levels predicting GF. Without the HR increase levels (Figure 2a, model A), communicative cue condition and reliability predicted GF respectively (communicative cue: estimate ± SE = 0.2279 ± 0.5132, *Z* = - 2.252, *p* =.0243; reliability: estimate ± SE = 0.1845 ± 0.4884, *Z* = −2.142, *p* =.0214). Both of communicative cue condition and reliability predicted HR increase levels (communicative cue: estimate ± SE = 0.0011 ± 0.0049, *Z* = −4.699, *p* <.001; reliability: estimate ± SE = 0.00085 ± 0.0039, *Z* = −4.6, *p* <.001). However, with HR increase levels, communicative cue condition and reliability did not show direct effects on GF (communicative cue: estimate ± SE = 0.2437 ± 0.1908, *Z* = −0.783, *p* =.434; reliability: estimate ± SE = 0.1965 ± 0.1864, *Z* = −0.949, *p* =.343). In other words, HR increase mediated between communicative cue condition or reliability and GF (Figure 2b, model B). Model B (619.47) showed a smaller Akaike Information Criterion (AIC) than model A (667.76), thus the mediation model was chosen as the better model.

**Figure 2.**
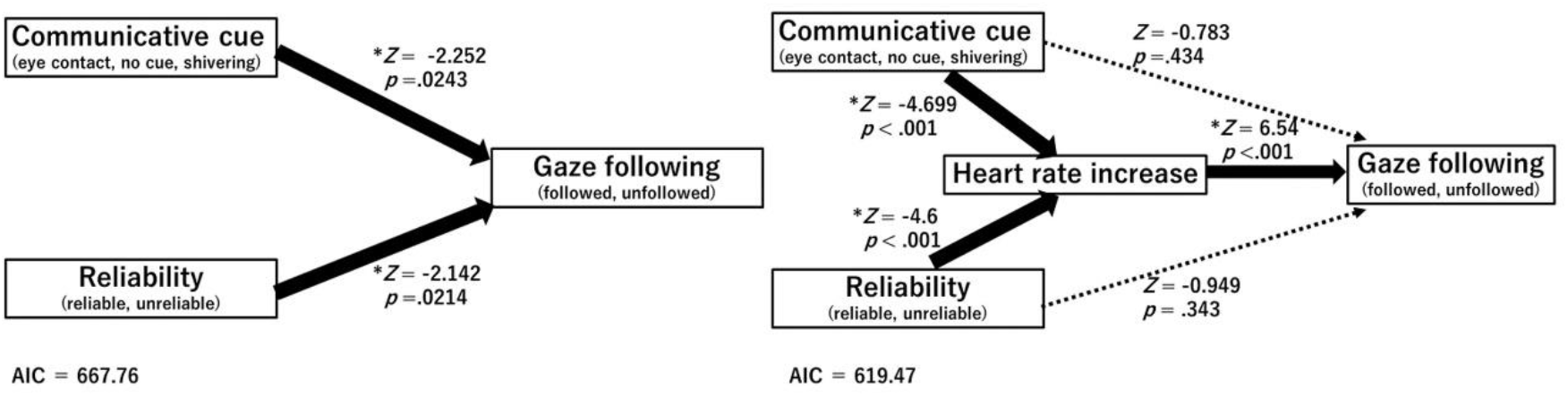
(a) Regression model predicting gaze following by communicative cue conditions (eye contact, no cue, shivering) and reliability (reliable, unreliable); (b) mediation model shows that heart rate increase mediates relationship between communicative cue conditions or reliability and gaze following.

## Discussion

The study revealed that the eye contact and reliability modulate the probability of infant GF and infant heart rates respectively, suggesting each contextual effect on GF is independent. While each contextual factor affected infant GF independently, the result of the mediation analysis showed that the increase in heart rate fully mediated both effects of the eye contact and reliability on GF. These results are in line with our hypothesis that the various social context is integrated as an action value, and the reward expectation is based on the calculated action value and indexed by the heart rate [26, 27], then modulates infant GF.

We replicated that eye contact or reliability facilitate GF, and showed that these facilitative effects are independent. We considered that we did not find the interaction due to ceiling effects. Social behaviour is not uniform and humans do not always respond in the same way to their social partners. Thus, the probability of GF which is manipulated by contextual factors may have a ceiling. As we hypothesized, it is suggested that contextual factors modulate the probability of GF, not always inducing the same behaviours.

The results are consistent with our hypothesis that the physiological arousal before GF behaviour is an index of reward expectations, as the result of action value calculation. Convergently, previous adult studies have indicated that reward expectations induce high physiological arousal [26, 27]. Also, it has been shown that reward predictive cues can enhance physiological arousal in infants and the physiological response to reward-associated cues predicts the success of associative learning [25]. The ostensive signals have been described as cues which convey communicative intent to the social partner [3], which would have activated reward system as other communication acts such as engaging with speech [37] and looking to the same object with the other person (joint attention) [38]. Thus, infants may be able to expect a later social reward from ostensive signals which promote GF. Also, in the current study, infants observed gaze cueing situations whether the female gaze direction is reward predictive or not (or the reliability of the actor). Therefore, in a socially engaging context, infants may calculate an action value of social interactions by integrating multiple social contextual cues (immediate visual context and knowledge), and the reward expectations during facing the reliable looker would then increase infant physiological arousal. Because the physiological state is known to reflect the calculated action value or expected reward in the social context [25], the results speculatively suggest that the contextual effects on the probability of infant GF, or infants’ behavioural choice whether or not to follow gaze, was mediated by the increase of heart rate.

We further argue that the observed increase of heart rate before GF is driven by the representation of the reward expectations, based on the abundance of neuroimaging studies suggesting the link between the two. The reward/affective system is anatomically close to the insula, the anterior cingulate cortex (ACC), and the hypothalamus [39]. These regions play a critical role to modulate sympathetic activities, in other words, they modulate physiological states [40–42]. Based on the simultaneous measurement of brain activity and physiological states during decision making, it has been reported that the brain activation network among the reward/affective system, the insula, and the ACC correlates increase of physiological states [43, 44]. To further examine if the contextual modulation of infants’ heart rates (which then mediated GF) is based on the reward expectation, future studies will need to examine whether infants’ heart rates are correlated with activations in the brain reward system and heart rates can index reward expectations in general.

This study is in line with the hypothesis that the action value calculations underlie adaptive modulation of social behaviour for relevant social contexts in human infants. It has been suggested that animals calculate action values through the integration of external sensory signals with current internal states, and these decisions ideally lead to optimal behavioural choices even in a situation requiring immediate responses such as a situation encountering a predator [45]. In humans, it has been reported that stimuli and actions that are uniquely encountered in social interactions can reinforce behaviour through neural mechanisms that are similar to those underlying non-social reinforcement with money, suggesting that humans calculate expected social reward in social interactions [46]. Social interaction and communication provide crucial opportunities to learn about the external environment for human infants. For example, following other’s gaze direction plays an important role in language learning [47, 48] and detecting potential threat sources [49]. Thus, infants’ GF behaviour may be optimized for social learning or avoiding potential threats in each social context.

A remaining question is how GF behaviour is optimized throughout the course of early development. Cognitive theories of GF such as Natural Pedagogy [3] and Perceptual Narrowing account of GF emergence [50] can account for the development of GF behaviour within the first year of birth. These theories explain behavioural bias to follow gaze in specific situations and how infant GF become attuned. However, these theories fail to provide a more general account of contextual modulation of infant GF, beyond the specific context modelled in each theory. For example, it has been reported that 20-month-olds show GF equally in situations with or without ostensive cues [51]. Also, 12-18-month-olds followed their caregivers’ and strangers’ gaze equally [52]. It is considered that contextual cues become less effective on GF after the first year of life. A computational study suggested that infants tend to show GF regardless of contextual factors after they have learned the action value of GF [53]. Thus, contextual cues might have a higher weight in affecting reward expectations and GF in the earlier stage of development, when infants have not yet learned the action value of GF behaviour.

During this early stage, contextual cues such as ostensive cues may help infants to disambiguate the context and expect rewards of social interactions, which then facilitate infant GF. Infants could then update the assignment of action value of GF through the extensive experience of social communication through the development. This perspective could explain the inconsistencies within the literatures on infant GF, especially why specific contextual cues modulate GF early in the development but then they start to follow gaze in a wider context in later development, not requiring scaffolding through additional contextual cues.

Future studies should also address what is rewarding for infants in social interactions. On one hand, Tomasello [54] argued that social engagement such as joint attention is rewarding for infants due to the evolution of collaborative activities and sharing, not by the information richness. On the other hand, adult’s social signals could be rewarding because it is informative, as it has been shown that informative value can be a reinforcement value. Humans preferentially choose options that will provide more information when making decisions [55], and acquiring information enhance brain activations in the reward system modulating learning performance [56]. Action value to acquire information can be considered as a reinforcement value for humans and it modulates behavioural decision making. In GF situations, there could be a mixture of two rewards: the social reward of joint attention and the reward of acquiring information about the environment. Future studies could further examine how infants assign reward value to adult’s social signals in GF situations, and in a wider range of interactive social contexts.

GF studies in experimental settings have mainly focused on the functional explanation of particular experimental manipulation of available contextual cues, such as the presence of attention-grabbing actions [29], the facial familiarity of interacting partners [31], the contingent reactivity [57], and the experience of social interaction before GF task [58]. In the experimental settings of GF situations our study is based on [7, 8, 28], typically only two objects are presented and GF occurs with a probability of 50% applying the definition of the first fixation to a target object. While in a naturalistic context, the surrounding environment is more complex with many objects, and infants show a wider repertoire of behaviour than in a screen-based experiment, which greatly reduces the probabilities of infants’ GF in each event far below 50% [23, 59]. Thus, we need to be cautious not to simply generalise our findings on a screen-based eye-tracking study to a wide range of ‘GF’ studies, many of which are assessed in more naturalistic and complex contexts. It is necessary to examine how contextual factors modulate infants’ GF in naturalistic situations, before such generalisation can be made.

Infant GF can be hypothesised as a result of the value-driven decision-making process, which enables infants to actively select when, and from whom, to learn during social communication.

## Ethics

The experimental protocol was approved by the Research Ethics Review Board, Department of Psychology Kyoto University, Japan, and the Research Ethics Review, Board, Department of Psychology Doshisha University, Japan. The parents of all participants provided written informed consent before their infants took part in this study.

## Data accessibility

Data used in the analysis are available in the electronic supplementary material.

## Acknowledgments

We appreciate the cooperation of all families that agreed to participate in this study. We thank Hiroki Yamamoto for his help for discussions. We would also like to thank the anonymous reviewers and colleagues who have provided us useful feedback or helped to conduct experiments. This work was supported by MEXT Promotion of Distinctive Joint Research Centre Program Grant Number JPMXP0619217850.

## Author contributions

M.I. and A.S. developed the study concept. M.I. conducted experiments and data analysis. M.K. worked for the ethical approving procedure and experimental settings. S.I. supervised this study. All authors approved the hypothesis, methods, and discussed the results.

