## Supplemental Materials for "Affective arousal explains infant gaze following under various social context"

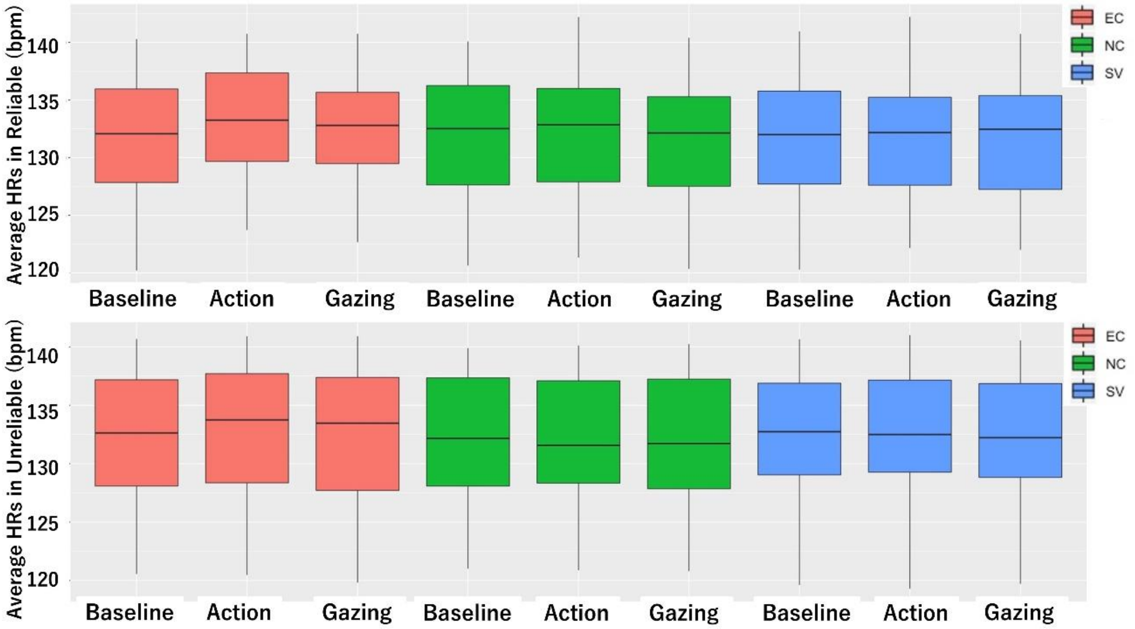


Figure S1. Mean heart rate levels during each phase for each condition. The x-axis depicts the video phase and the y-axis depicts the beats per minute of heart rate.

### Additional analysis

#### Attention to the face area

To confirm that infants equally learned the actor's face in gaze-cueing situations, we compared total fixation time to the face between reliable and unreliable using a t-test. There was no difference in total fixation time to the face area between reliable ($M$ = 1.23s) and unreliable ($M$ = 1.20s). Infants equally looked at the face before the gaze following task.

In the gaze following task, because it is possible that attention to the model's face affected infant HR and gaze following, we examined the infant's attention to the model's face across conditions in all three phases. We conducted an ANOVA using gaze time at the model's face as the dependent variable across all the conditions for each phase (two levels of reliability: reliable, unreliable; three levels of condition: EC, NC, SV; three-phase levels: Baseline, Action, Gazing). There were no differences between any conditions or phases.

#### Learning effects over the trials

Infants engaged in two blocks of the experimental procedure; therefore, repeated trials could affect infant learning during the experiment. To examine potential learning effects over the trials, we compared gaze following frequency between the first block and the second block of the experiment across all conditions (two levels of reliability: reliable, unreliable; three levels of condition: EC, NC, SV) using chi-square test. There was no significant difference between blocks ($\chi2(5) = 4.645$, $p$ =.461), thus there were no learning effects of repeated trials.
